# Loss of Grp170 results in catastrophic disruption of endoplasmic reticulum functions

**DOI:** 10.1101/2023.10.19.563191

**Authors:** Melissa J. Mann, Chris Melendez-Suchi, Maria Sukhoplyasova, Ashley R. Flory, Mary Carson Irvine, Anuradha R. Iyer, Hannah Vorndran, Christopher J. Guerriero, Jeffrey L. Brodsky, Linda M. Hendershot, Teresa M. Buck

## Abstract

GRP170, a product of the *Hyou1* gene, is required for mouse embryonic development, and its ablation in kidney nephrons leads to renal failure. Unlike most chaperones, GRP170 is the lone member of its chaperone family in the ER lumen. However, the cellular requirement for GRP170, which both binds non-native proteins and acts as nucleotide exchange factor for BiP, is poorly understood. Here, we report on the isolation of embryonic fibroblasts from mice in which LoxP sites were engineered in the *Hyou1* loci (*Hyou1^LoxP/LoxP^*). A doxycycline-regulated Cre recombinase was also stably introduced into these cells. Induction of Cre resulted in excision of *Hyou1* and depletion of Grp170 protein, culminating in apoptotic cell death. As Grp170 levels fell we observed increased steady-state binding of BiP to a client, slowed degradation of a misfolded BiP substrate, and BiP accumulation in NP40-insoluble fractions. Consistent with disrupted BiP functions, we observed reactivation of BiP storage pools and induction of the unfolded protein response (UPR) in futile attempts to provide compensatory increases in ER chaperones and folding enzymes. Together, these results provide insights into the cellular consequences of controlled Grp170 loss and insights into mutations in the *Hyou1* locus and human disease.

## Introduction

Approximately one-third of the proteins encoded by the human genome will enter the endoplasmic reticulum (ER) co- or post-translationally (Dancourt and Barlowe, 2010) where they will undergo a variety of modifications, fold, and assemble into their native form (Braakman and Bulleid, 2011),(Phillips and Miller, 2021). This process is monitored by a dedicated ER quality control system (ERQC) that ensures only properly matured proteins are transported to the Golgi and to points further along the secretory pathway (Adams et al., 2019),(Pobre et al., 2019). To meet these demands, ERQC is comprised of a large complement of resident molecular chaperones, their co-factors, and two groups of folding enzymes, which bind unfolded polypeptide chains as they enter the ER lumen. Together they protect vulnerable hydrophobic or aggregation prone regions on nascent proteins until they are buried upon folding, as well as to prevent the transport of immature intermediates. Equally important to the fidelity of the secretome is the need to identify proteins that fail to complete folding and/or assembly and target them for degradation by the proteasome (Vembar and Brodsky, 2008),(Krshnan et al., 2022),(Christianson et al., 2023). This ER-associated degradation (ERAD) pathway relies on some of the same chaperones and folding enzymes that assist protein folding and assembly in the ER to deliver misfolded proteins to a retrotranslocon complex for extraction to the cytosol (Needham et al., 2019),(Wu and Rapoport, 2021). Failure to match the demands of the biosynthetic load of the ER triggers the unfolded protein response (UPR) that strives to restore homeostasis, but if unsuccessful, UPR activation results in cell death (Walter and Ron, 2011),(Hetz et al., 2020).

Many of the nascent proteins that enter the ER will ultimately reside in organelles of the secretory pathway, or they will be expressed at the cell surface or secreted into the extracellular space. This diverse polypeptide ensemble is uniquely modified by disulfide bonds and is N-linked glycosylated, which serves in part to stabilize them in spaces that are chaperone-depleted. Over 20 ER-resident protein disulfide isomerases (PDIs) assist in the formation, reduction, or isomerization of disulfide bonds (Appenzeller-Herzog and Ellgaard, 2008),(Robinson and Bulleid, 2020), and two lectin chaperones and their co-factors assist in the maturation of glycoproteins (Molinari and Hebert, 2015). Disruption of individual members of these components of the ERQC machinery is largely tolerated due to redundancy among family members. Yet, in addition to the unique modifications to which proteins in the ER are subject, more general aspects of protein folding in the ER follow the same mechanisms that guide the transition of nascent polypeptide chains to their native conformations in the cytosol. In keeping with this view, the ER also possesses multiple members of both the cyclophilin and FKBP prolyl-peptidyl isomerase (PPIase) families, which catalyze the *cis-trans* isomerization of peptide bonds N-terminal to specific proline residues (Schiene-Fischer, 2015).

In contrast to the redundancy provided by the chaperone/folding families with multiple members, the mammalian ER possesses a single cognate of the Hsp70 (BiP/GRP78), Hsp90 (GRP94), and Hsp110 (GRP170) chaperone families. BiP, the first chaperone discovered in the ER, was identified through its association with unassembled immunoglobulin heavy chains (Bole et al., 1986; Haas and Wabl, 1984). BiP recognizes short hydrophobic polypeptide stretches on nascent proteins (Blond-Elguindi et al., 1993),(Flynn et al., 1991) that are ultimately buried in the native state, providing its specificity for unfolded proteins. Genetic disruption of BiP in mice is embryonic lethal at day e3.5 (Luo et al., 2006), and the AB5 subtilase cytotoxin kills cells by cleaving BiP between the client binding domain and regulatory nucleotide binding domain (Paton et al., 2006), findings that reveal the essential nature of this chaperone. Although BiP is the only Hsp70 family member in the ER, it is assisted by one of eight DnaJ family co-chaperones. Four of these, ERdj3-6, can bind a variety of clients and transfer them to BiP, which allows these chaperones to contribute to diverse and even opposing ER functions (Pobre et al., 2019). At the cellular level, deletion of individual members of the ERdj family is tolerated and single knock-out cells are viable (Amin-Wetzel et al., 2017).

Like BiP, GRP94—the lone Hsp90 family member in the ER—is required for mouse embryonic development, with null mice dying between day e6.5 - e7.5 (Wanderling et al., 2007). Similar to cytosolic Hsp90, GRP94 has a more restricted clientele, which likely accounts for the ability to disable the gene in some tissues but not others. For instance, a mouse pre-B cell line was identified that lacked GRP94 (Randow and Seed, 2001), and embryonic stem cells can be isolated from a constitutive knock-out mouse, which will differentiate into adipocytes, hepatocytes, and neurons, but not into myocytes (Wanderling et al., 2007). Similarly, tissue-specific expression of Cre used to delete the *GRP94*-encoding gene is readily tolerated in the mammary gland (Zhu et al., 2014), but Cre expression in myeloid precursors (Luo et al., 2011), hepatocytes (Poirier et al., 2015), or pancreatic β islet cells (Kim et al., 2018) alters differentiation or adversely affects function.

GRP170, which belongs to the family of large Hsp70 proteins that includes cytosolic Hsp110, is the lone ER member of this chaperone family. GRP170 acts as a nucleotide exchange factor (NEF) for BiP to stimulate its release from clients (Steel et al., 2004),(Weitzmann et al., 2006). In addition to its NEF activity, GRP170 acts as a molecular chaperone that binds aggregation-prone sequences in non-native proteins (Behnke and Hendershot, 2014; Behnke et al., 2016; Park et al., 2003). GRP170 has also been shown to regulate the maturation of the epithelial sodium channel (ENaC) as it matures in the ER and participates in the ERAD of unassembled subunits (Buck et al., 2013). Attempts to produce Grp170 null mice were unsuccessful (Kitao et al., 2001), revealing an essential role for this chaperone during mouse development, despite the presence of a structurally-unrelated, second NEF in the ER lumen, Sil1 (Chung et al., 2002). These data are consistent with unique functions of the two ER NEFs and is further supported by the identification of a human disease directly linked to SIL1 mutations (Anttonen et al., 2005),(Senderek et al., 2005). Additional support for the unique activities of these two ER NEFs was provided by the generation of a Cre-inducible nephron-specific mouse strain in which LoxP sites were engineered in the *Hyou1* loci (*Hyou1^LoxP/LoxP^*). Regulated expression of Cre recombinase in kidney nephrons led to rapid excision of this gene, resulting in mice with hallmarks renal cell injury of acute kidney injury (Porter et al., 2022). Yet, the cellular and molecular mechanisms underlying renal cell damage were ill-defined.

To this end, we created an experimentally tractable system to characterize the role of Grp170 in regulating various aspects of ER functions and response to stress. Specifically, mouse embryonic fibroblasts (MEFs) were obtained from *Hyou1^LoxP/LoP^*and doxycycline-regulated Cre recombinase was introduced. Single cell clones were isolated that rapidly and reproducibly resulted in complete excision of the *Hyou1* gene upon doxycycline treatment. Even though *Sil1* was induced, the depletion of murine Grp170 led to wide-spread disruption of ER homeostasis and activation of the UPR, culminating in apoptotic cell death. These data further support unique activities of the two ER lumenal NEFs, highlight previously uncharacterized features of Grp170 function in the ER, and describe a tool to better define new Grp170 attributes.

## Results

### Isolation of Hyou1^LoxP/LoxP^ mouse embryonic fibroblasts and creation of an inducible Cre construct

Fibroblasts were isolated from 12.5 d embryos generated from *Hyou1*^LoxP/LoxP^ female mice that has been crossed with *Hyou1*^LoxP/LoxP^ males, and cultured cells were transformed with SV40. In an attempt to produce stable Grp170 null MEFs, the cells were transduced with lentivirus expressing GFP and Cre recombinase under control of a constitutive SV40 promoter (Qin et al., 2010). However, two separate viral transductions with a multiplicity of infection (MOI) of 2 resulted in no GFP positive cells (data not shown), suggesting that MEFs might be unable to tolerate loss of Grp170. Thus, a lentivirus backbone was engineered with a doxycycline-inducible Cre recombinase separated from mCherry with a P2A ribosome cleavage peptide (Supplemental Figure 1A). The construct was first tested in 293T cells, and after 2 days, GFP and mCherry expression were monitored by FACS. In the absence of doxycycline, GFP was readily detected with only a minor population of cells that also expressed mCherry (Supplemental Figure 1B), whereas doxycycline addition resulted in linear expression of mCherry corresponding to GFP levels. Cell lysates were then examined for Cre expression by western blotting, which revealed that mCherry fluorescence faithfully reported on Cre recombinase expression (Supplemental Figure 1C).

### Continuous expression of Cre recombinase is selected against in Hyou^LoxP/LoxP^ MEFs

We next introduced the Cre-modified lentivirus into *Hyou*^LoxP/LoxP^ MEFs and sorted the cells to isolate a bulk culture of GFP^+^ and mCherry^−^ cells. Although this bulk culture was selected based on GFP expression, we observed a population of cells that were no longer GFP^+^ as the sorted cells were expanded (Figure 1A, day 0). The cultured cells were treated with doxycycline throughout the time course to induce Cre recombinase expression, and both GFP and mCherry fluorescence were monitored at various time points thereafter (Figure 1A). Cre recombinase induction and its effect on Grp170 expression were directly examined by RT-PCR and western blot. After two days of doxycycline treatment, ∼50% of the cells co-expressed both fluorescent proteins. However, several changes were observed over time. First, the population of cells expressing mCherry (and presumably Cre recombinase) dropped precipitously until they virtually vanished. Second, a little over 50% of the cells at later time points expressed only GFP, indicating that cells in which mCherry was not induced by doxycycline were being selected. Third, the minor population of cells that did not express GFP at the beginning of the time course had expanded until they comprised 50% of the culture by day 11. This suggested that extinguishing Grp170 expression was toxic to MEFs.

**Figure 1.**
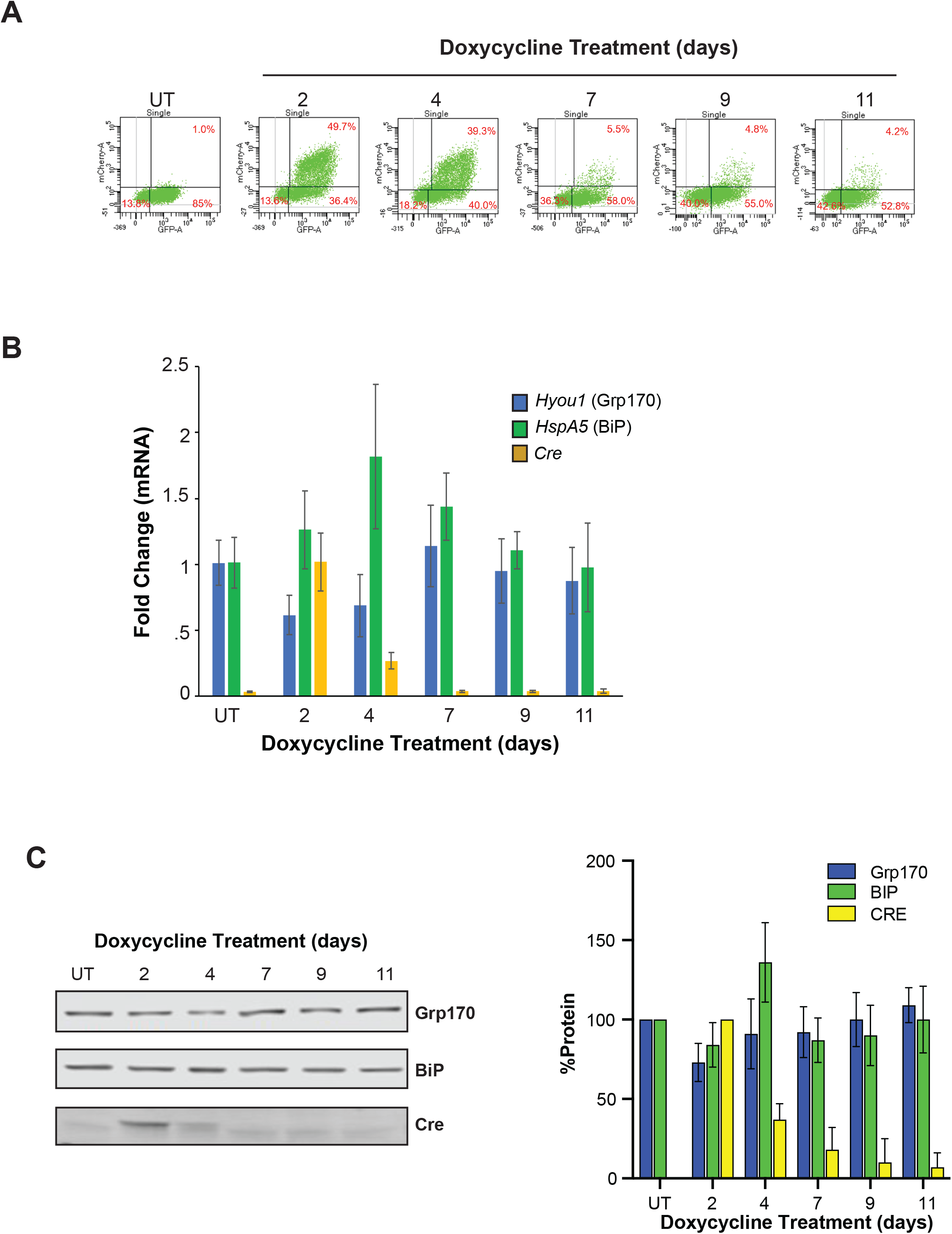
*Hyou1^fl/fl^*MEF cells expressing Cre recombinase are rapidly out-competed by cells that do not. **A.** GFP and mCherry expression were measured by FACScan at the indicated days after incubation with doxycycline. **B.** The indicated transcripts were quantified by RT-PCR after doxycline treatment to induce Cre synthesis. **C.** Corresponding protein expression of the transcripts measured in **B** were determined by western blotting. The signals were quantified and graphically displayed as a percent of their value in untreated cells (n=3). Data presented represent the mean +/−SD, *p<0.05, **p<0.01, ***p<0.001.

In keeping with the pattern observed for mCherry expression, Cre mRNA and protein were readily observed after two days of doxycycline treatment. Cre expression was dramatically reduced by day 4 of doxycycline treatment and was undetectable thereafter (Figure 1B and C). Correspondingly, after two days of treatment, *grp170* mRNA and protein decreased to ∼60% and ∼80% respectively, with similar levels on day 4. Since only ∼50% of cells expressed mCherry/Cre, we surmised that Grp170 protein levels in the Cre positive cells were likely ∼30 - 40% of that observed in untreated cells on these respective days. Consistent with the nearly complete loss of cells expressing mCherry/Cre from the bulk population, by day 7 Grp170 levels were the same as controls. We also monitored BiP levels since BiP shares clients with GRP170 (Behnke and Hendershot, 2014) and—as noted above—is the target of GRP170’s nucleotide exchange activity (Steel et al., 2004; Weitzmann et al., 2006). The expression of BiP transcripts and protein increased as Grp170 levels declined and peaked on day 4. As Cre^+^ cells disappeared and Grp170 levels normalized, BiP transcripts also returned to baseline, although BiP protein levels were slower to decline (Figure 1B and C), most likely due to its long half-life (Wang et al., 2017).

### Isolation of single cell Hyou^LoxP/LoxP^ clones expressing inducible Cre recombinase

Because we were unable to isolate a bulk culture of Grp170 null MEFs, we next isolated single cell clones in an effort to obtain cells with tight suppression of Cre recombinase in the absence of doxycycline, and to determine if complete ablation of Grp170 expression was possible with doxycycline treatment. To do so, we isolated the brightest ∼15% of GFP^+^ cells from the bulk culture and chose 88 single cell clones to expand and characterize. An aliquot of each clone was treated with doxycycline and examined for GFP and mCherry expression. Of these, only five clones were >90% positive for both GFP and mCherry (Table 1). These results were obtained despite each clone arising from a single cell, underscoring the enormous selective pressure against expressing Cre recombinase and thus eliminating Grp170 expression. Nevertheless, two of these clones were >99% positive for both fluorescent proteins (clone A10 and E8) and were used for further analyses.

To begin to characterize the A10 and E8 clones, the kinetics and magnitude of doxycycline-induced Grp170 deletion was examined using two sets of primers that identified the intact gene and one set that detected the Cre-recombined, deleted gene (Supplemental Figure 2A, B). Within 18 hrs, the intact gene was no longer detected, suggesting that disruption of the Grp170-encoding alleles was complete. When the primer set that detected the gene deletion was used, we observed an unexpected signal in the absence of doxycycline, although this signal increased over time with doxycycline treatment (Supplemental Figure 2A). This result was also observed in the other three clones with high inducible mCherry expression (data not shown). To further characterize these clones, we performed fluorescence *in situ* hybridization (FISH) analysis on the A10 and E8 clones before and after doxycycline treatment. Both lines were polyploid, which is a consequence of SV40-transformation (Ray et al., 1990). In the A10 clone, 98.5% of the cells had three copies of the region surrounding the *Hyou1* locus, as detected with a probe for the *Hyou1* telomeric flanking region (red), only two of which still contained the *Hyou1* gene (green) (Supplemental Figure 2C, untreated). The loss of *Hyou1* from the third copy explained the signal with the primer pair for a rearranged allele. In the E8 clone, 97% of the cells had four copies of this locus. Of these, 58% had one *Hyou1* allele deleted and 42% had two alleles deleted. Thus, all the cells in both clones retained at least two copies of the *Hyou1* gene. Nevertheless, after Cre recombinase induction with doxycycline, we observed deletion of the gene in 100% of the cells in both clones, regardless of how many copies of the gene they possessed (Supplemental Figure 2C).

### Grp170 is an essential protein

The levels of basal *Grp170* transcripts and protein in these two clones were compared to the non-transformed MEFs and to the SV40-transformed MEFs in which the dox-inducible Cre system had not been introduced. TheSV40-transformed MEFs expressed about twice as much *Grp170* mRNA compared to the non-transformed MEFs (Supplemental Figure 2D). In keeping with increased transcript levels, the bulk culture of transformed MEFs possessed 3-4 copies of the *Hyou1* locus compared to the non-transformed cells, which had 2 copies of this gene (see above). It was noteworthy, however, that Grp170 protein levels remained near that of the non-transformed cells, although transcript levels were much higher. These results might be consistent with an auto-regulatory feed-back system, which has been reported for BiP and cytosolic Hsp90 (Gülow et al., 2002),(Cheng et al., 2010). As shown in Supplemental Figure 2D, the A10 clone expressed *Grp170* transcript levels that were similar to that of the non-transformed MEFs, whereas the E8 cells had about half the amount of *Grp170* transcript, indicating that this reduction in protein level did not adversely affect MEF growth.

Doxycycline treatment of the A10 and E8 clones resulted in a significant reduction in *Grp170* transcripts, which was apparent at day 3 of doxycycline treatment (Figure 2A). Protein levels began decreasing within 1 day and dropped precipitously to <5% by days 4 and 6 in the A10 and E8 clones, respectively (Figure 2B). Unlike the bulk cultures (see above), there was no “rebound” expression of Grp170 protein during this period. Moreover, as Grp170 protein levels diminished, cells began lifting off the dishes and were unable to exclude Trypan Blue (data not shown). In accordance with these data, viability of both cell lines began to decrease within 1 day of doxycycline treatment and continued to fall as Grp170 protein levels decreased (Figure 2B). Cell cycle analyses revealed that cell death was not accompanied by arrest at any particular phase of the cell cycle (data not shown). Together, in keeping with our inability to isolate stable clones that were deficient in Grp170 expression (see above), the cells in both the A10 and E8 clones died as Grp170 was progressively depleted, despite concurrent up-regulation of Sil1 (Figure 2D).

**Figure 2.**
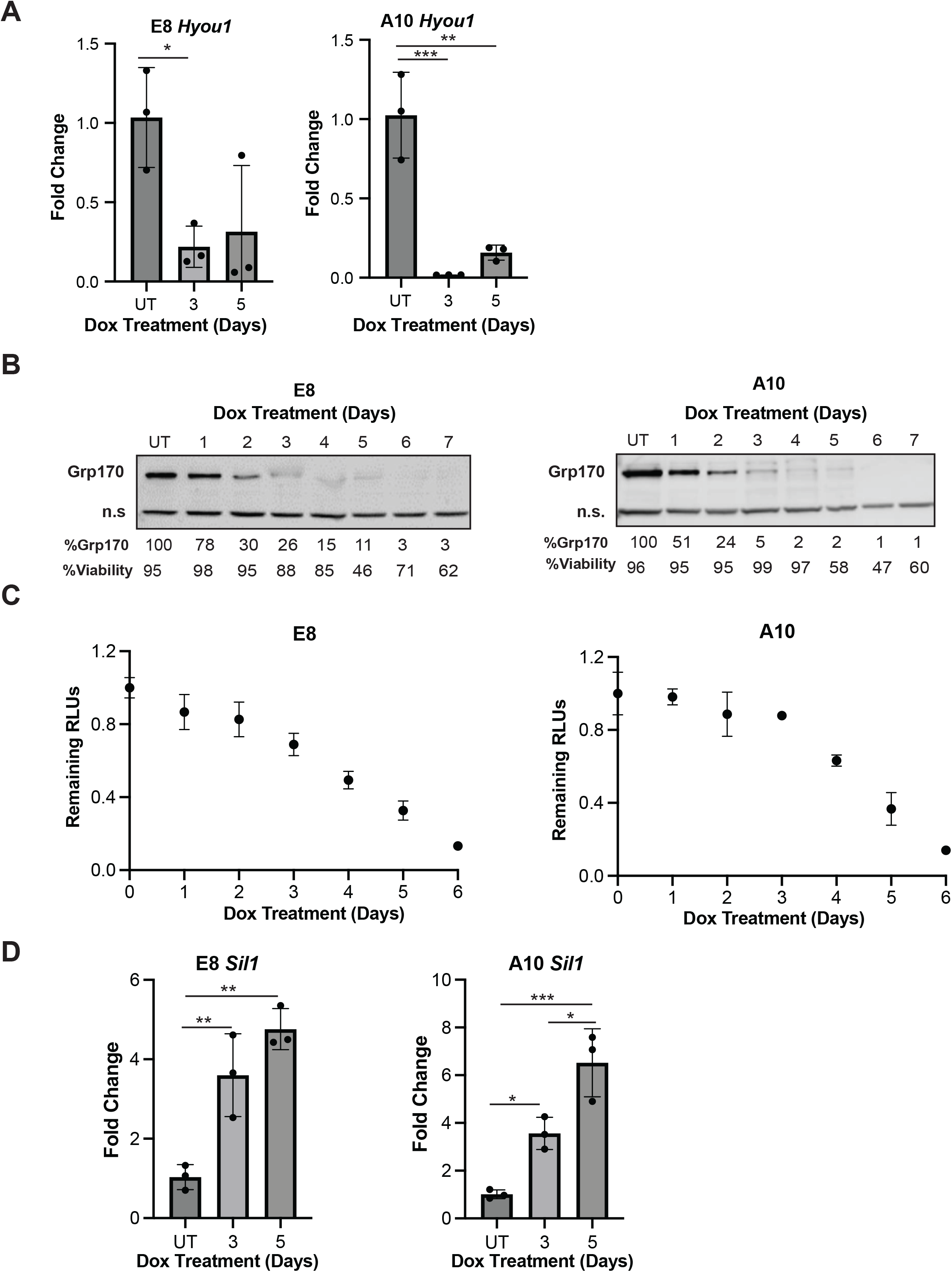
Inducible expression of Cre recombinase in single cell clones of *Hyou1^fl/fl^* MEFs leads to loss of Grp170, which is accompanied by cell death. **A.** The expression of *Hyou1* transcripts were measured in the E8 and A10 clones after Cre induction (n=3). **B.** The resulting effect on GrpRP170 protein expression was assessed by western blot with levels relative to untreated cells displayed below each time point. n.s. indicates an non-specific band. **C.** Cell viability was determined each day after Cre induction using a Cell Titer-Glo assay. Signals were expressed relative to that in untreated cells (n=3). **D.** Transcripts for *Sil1*, the other ER NEF, were quantified by RT-PCR after loss of *Grp170*.

### Loss of Grp170 results in dysregulated BiP function

We next examined a number of ER functions to understand why cells die in the absence of Grp170. We first focused on BiP, given that Grp170 is a NEF for BiP, Grp170 shares a number of clients with BiP, and all of BiP’s functions—other than its contribution to ER calcium stores (Lievremont et al., 1997)—are adenosine nucleotide dependent (Behnke et al., 2014).

Under homeostatic conditions, a pool of BiP is AMPylated (Chambers et al., 2012). This reversable modification provides an inactive reservoir of BiP that can be rapidly reactivated by ER stress (Perera and Ron, 2023) before the UPR can restore homeostasis by transcriptionally and translationally up-regulating a suite of resident ER chaperones and folding enzymes (Ron and Walter, 2007). In both the A10 and E8 clone, loss of Grp170 resulted in a rapid disappearance of AMPylated BiP (Figure 3A-B). Activating this pool of BiP suggests an increased demand for BiP. Therefore, we next explored BiP’s best understood function, the binding to an unfolded client. Because preliminary transient transfection experiments with GFP uncovered a significantly higher percent of the E8 clones expressed GFP than the A10 clone, we focused on the E8 clone to obtain quantitative data, but where indicated results were verified with the A10 clone.

**Figure 3.**
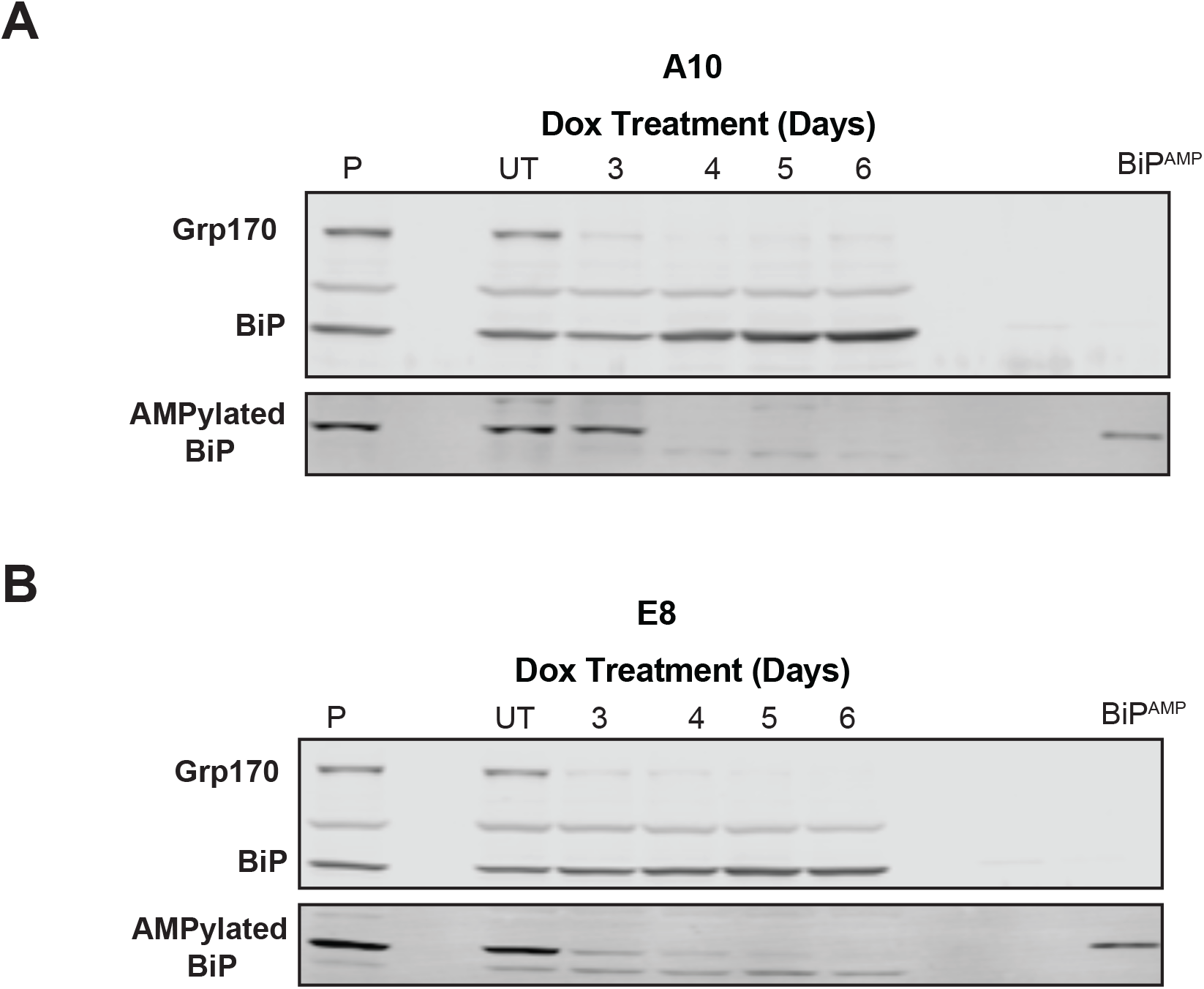
As Grp170 levels fall, AMPylated pools of BiP are reactivated to increase the pool of functional BiP. **A.** Cells from the A10 clone were incubated with doxycycline for the indicated days and cell lysates were prepared for western blotting. The top panel portrays a membrane blotted with a combination of anti-Grp170 and anti-BiP, whereas the bottom panel is from a membrane blotted with a monoclonal specific for AMPylated BiP. An aliquot of recombinant AMPylated BiP was loaded on the right of the gel. The parental clone (P) was loaded to the left. **B.** Cells from the E8 clone were analyzed as in **A**.

The non-secreted immunoglobulin (Ig) light chain variant, NS1 κ light chain, is a well-studied BiP client whose folding requires assembly with Ig heavy chains (Ma et al., 1990),(Skowronek et al., 1998). The interaction of BiP with unassembled NS1 κ light chains was determined by immunoprecipitation-coupled western blotting in cells treated with doxycycline for 5 days, which is sufficient to deplete ∼90% of Grp170 protein compared to untreated cells (Figure 2B). As shown in Figure 4A, we detected enhanced BiP association with NS1 under steady state conditions in the doxycycline-treated/Grp170-depleted cells. Because NS1 has a single ∼100 amino acid domain that folds poorly (Skowronek et al., 1998), this likely represents stabilization of binding as opposed to the binding of additional BiP molecules. Next, to determine if loss of Grp170 stabilized BiP binding to other clients, we examined the turnover of the Null Hong Kong (NHK) variant of α1-anti-trypsin. The retrotranslocation of this ERAD client to the cytosol for proteasomal degradation is dependent on BiP release (Inoue and Tsai, 2016). Consistent with defects in client release from BiP, pulse-chase experiments conducted on cells treated with doxycycline for 5 days revealed that NHK turnover via the ERAD pathway was significantly reduced (Figure 4B).

**Figure 4.**
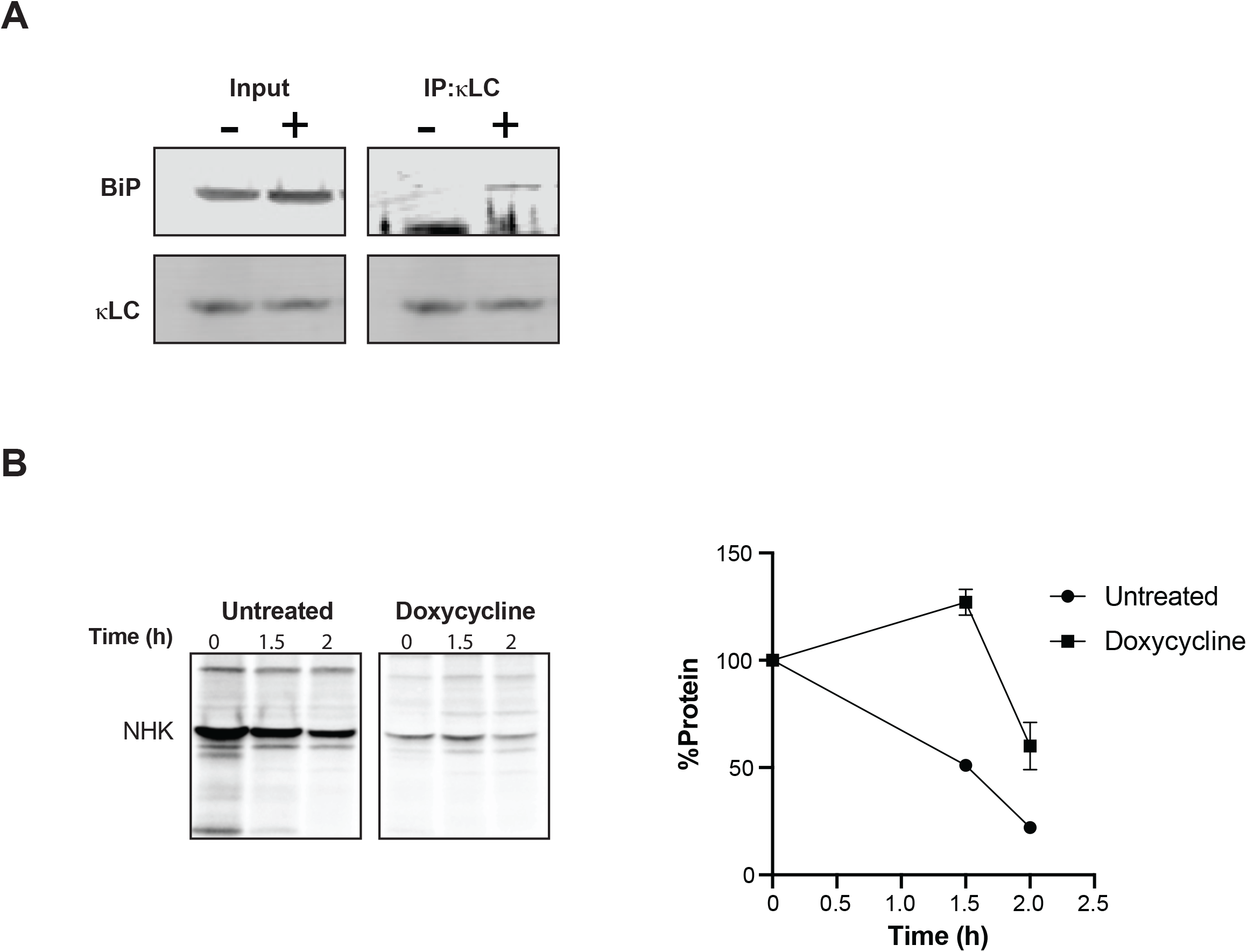
Loss of Grp170 increases steady-state BiP binding to unfolded clients and interferes with their degradation. **A.** Cells from the E8 clones were transfected with a vector encoding the NS1 LC four days after incubation with doxycycline and lysates were prepared the following day. Untreated cells served as a control. The mouse κ LC was immunoprecipitated from lysates, separated on reducing SDS-gels, transferred and blotted with anti-BiP or anti-κ. **B.** E8 cells that had been treated with doxycycline or left untreated were transfected with the NHK variant of α1AT. A day later cells were pulsed with [35S] methionine and cysteine for 30 minutes prior to chasing for the indicated times. Lysates were prepared, immunoprecipitated with anti-α1AT and electrophoresed on reducing SDS-polyacrylamide gels. Signals were determined on a phosphorimager and represented as a percentage of that observed at the end of the chase (t=0). Data represent the mean+/−SD.

Unlike BiP, which binds short hydrophobic sequences in unfolded proteins, GRP170 prefers longer amino acid sequences in its clients that are rich in aromatic residues and have a propensity to induce the formation of protein aggregates (Behnke et al., 2016). Notably, the introduction of a GRP170 binding site into the unfolded domain of an immunoglobulin heavy chain resulted in the formation of insoluble aggregates in human 293T cells, even though BiP binds this domain (Hendershot et al., 1987). To determine if BiP might be trapped in insoluble aggregates when Grp170 was deleted in the MEFs, we induced Cre expression and resolved proteins in the lysates into detergent-soluble and -insoluble fractions 3-6 days after doxycycline treatment. As Grp170 levels fell, BiP began to accumulate in the detergent-insoluble fraction (Figure 5).

**Figure 5.**
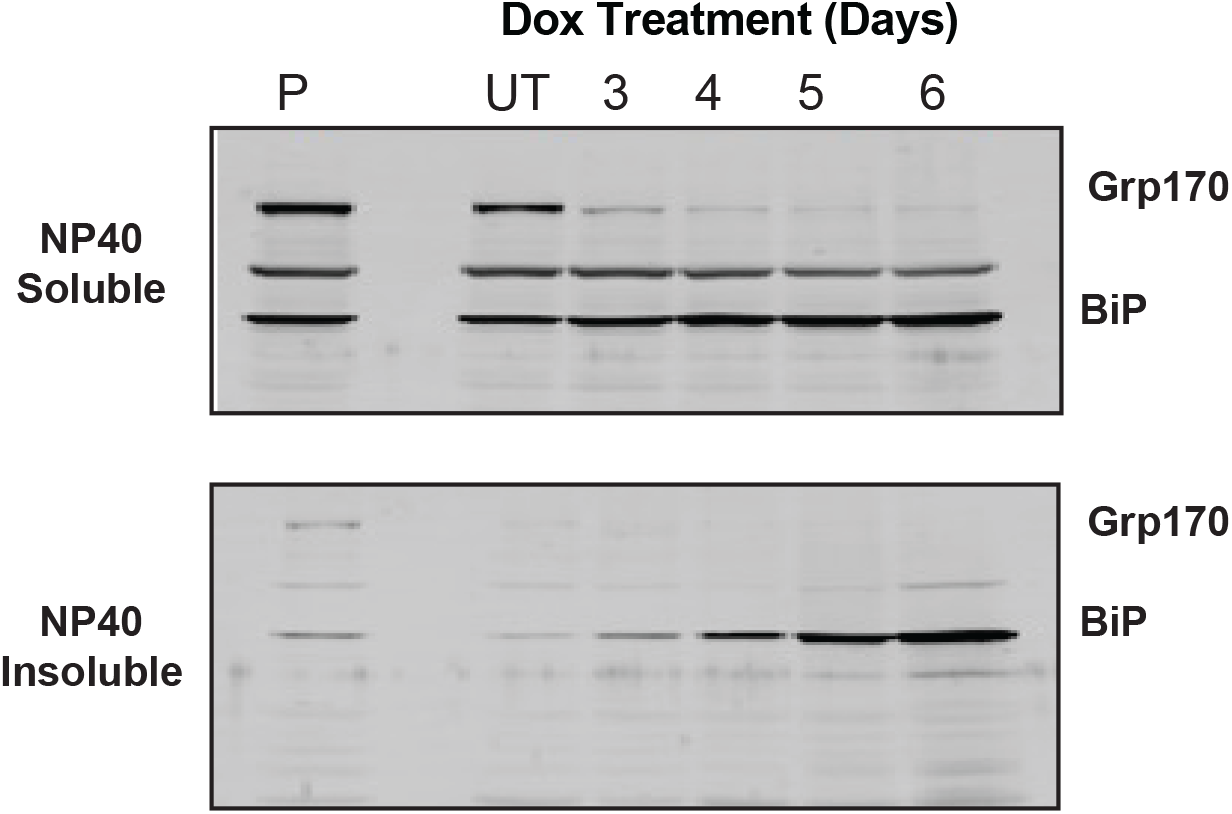
In the absence of Grp170, a portion of BiP fractionates with protein aggregates. Cre recombinase expression was induced in E8 cells for the indicated days. Cells were lysed with NP40 buffer and centrifuged to separate NP40 soluble and insoluble material. The insoluble material was extracted by heating in Laemmli buffer and sonication. The samples loaded on the NP-40 insoluble gel (bottom) corresponded to six times that loaded on the NP40 soluble gel (top). After electrophoresis and transfer, membranes were blotted for BiP and Grp170.

### Unresolved ER stress response culminates in apoptotic cell death

When BiP levels become insufficient to perform its various functions in maintaining ER homeostasis, it is recruited from the ER-localized UPR transducers, Ire1, ATF6 and PERK to aid in protein refolding/maturation or target non-native proteins for ERAD (Bertolotti et al., 2000),(Shen et al., 2002). This results in activation of the UPR transducers and signals a complex transcriptional and translational response to restore homeostasis, in part by up-regulating BiP (Walter and Ron, 2011),(Hetz et al., 2020). To further determine if Grp170 loss was adversely impacting BiP availability, both clones were examined for evidence of UPR activation (Figure 6A-B and Supplemental Figures 3-4). As described below, targets of each of the three arms of this ER stress response were up-regulated at both transcriptional (Figure 6A and S3) and translational (Figure 6B and Supplemental Figure 4) levels as Grp170 was depleted. Although there is significant cross-talk between the three downstream targets of the various transducers, the initial responses include: (1) activation of Ire1, thus triggering the splicing of the transcript encoding the *Xbp1* transcription factor (*sXbp1*), which increases BiP and Hrd1 levels; (2) ATF6 activation, similarly resulting in BiP up-regulation, and (3) activation of PERK, resulting in the phosphorylation (“p-eIF2α”) and inhibition of eIF2α translational factor activity, which then regulates ATF4 and CHOP. We found that the levels of these targets continued to increase through day 5 of doxycycline treatment, but by day 7 some of them decreased, with PERK transcriptional targets returning to basal levels. Interestingly, p-eIF2α protein levels remain high. Through a feedback loop, the translational inhibition caused by eIF2α phosphorylation drives synthesis of the Gadd34 phosphatase, which acts on p-eIF2α to restore translation (Harding et al., 2009). In keeping with prolonged eIF2α phosphorylation, lower levels of Gadd34 were evident at days 5 and 7 of doxycycline treatment (Figure 6C and Supplemental Figure 4). However, in spite of prolonged phosphorylation eIF2α, this pathway was no longer able to drive transcription of its targets ATF4 or CHOP at later time points.

**Figure 6.**
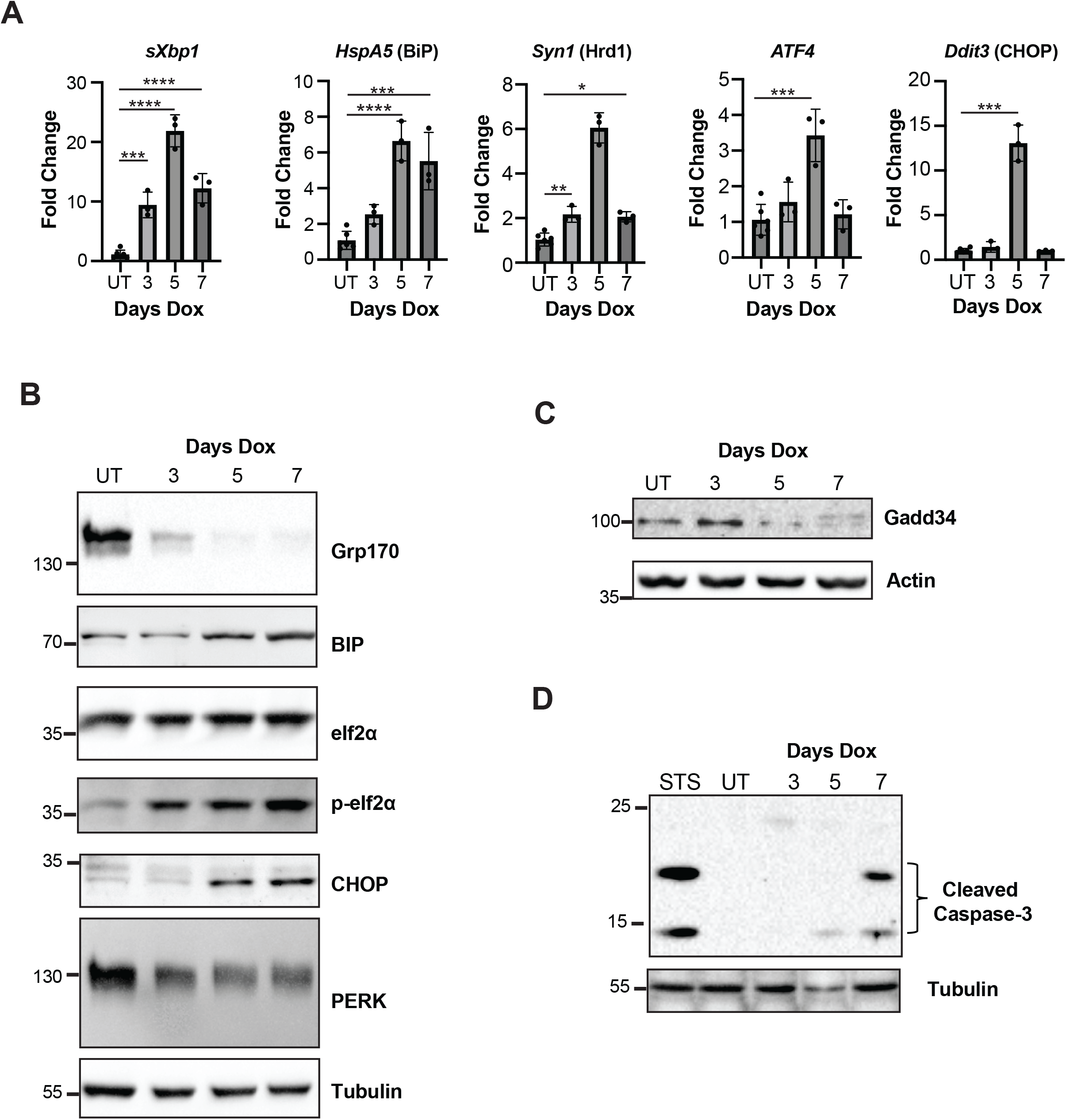
The unfolded protein response (UPR) is activated in the absence of Grp170. **A.** The expression of UPR target transcripts was measured in the E8 clones after Cre induction for the indicated days by qPCR (n=3), data represents the mean+/−SD; *p<0.05, **p<0.01, ***p<0.001, ****p<0.0001. **B.** Western blot analysis was also performed for UPR targets (**B**) Gadd34 (**C**), an apoptotic marker (**D**) or loading control (actin or tubulin). Representative images are shown.

In addition to a well-established role for CHOP in ER stress-associated cell death (Zinszner et al., 1998), a number of caspases are also activated by unresolved UPR activation (Iurlaro and Muñoz-Pinedo, 2016). One route by which apoptotic cell death occurs is via activation of caspase-3, downstream of Ire1 activation (Fribley et al., 2009). Consistent with the progressive onset of cell death observed in Figure 2B, caspase-3 cleavage was detected as early as day 5 post-doxycycline treatment and became significantly more prominent by day 7 (Figure 6D).

### Loss of Grp170 alters ER morphology

Sustained ER stress can disrupt ER morphology (Gardner et al., 2013),(Ng et al., 2000). To examine the effects of Grp170 loss on ER structure in MEFs, we stained fixed doxycycline-treated and control cells with an anti-KDEL antibody, which recognizes the ER retention sequence found on the C-terminus of resident ER soluble proteins (Munro and Pelham, 1987), and can be used to visualize the ER. As presented in Figure 7A, ER structure at days 5 and 7 post-doxycycline treatment appears more collapsed than the untreated or day 3 cells. Notably, the cells at later time-points also appear smaller, which was verified by quantifying the cytosolic GFP signal (Figure 7B). We next measured levels of Climp63, a cytoskeleton-linking membrane protein (Klopfenstein et al., 2001) that helps maintain ER morphology and is required for ER sheet formation (Shibata et al., 2010),(Gao et al., 2019). As Grp170 levels were depleted, Climp63 expression was unchanged at day 3 but was significantly reduced at days 5 and 7 (Figure 7C), consistent with a more collapsed ER. Decreased Climp63 expression may be due to reduced protein synthesis (see for example NHK signal in Figure 4B), which we attribute to sustained eIF2⍺ phosphorylation and reduced Gadd34 levels (Figure 6).

**Figure 7.**
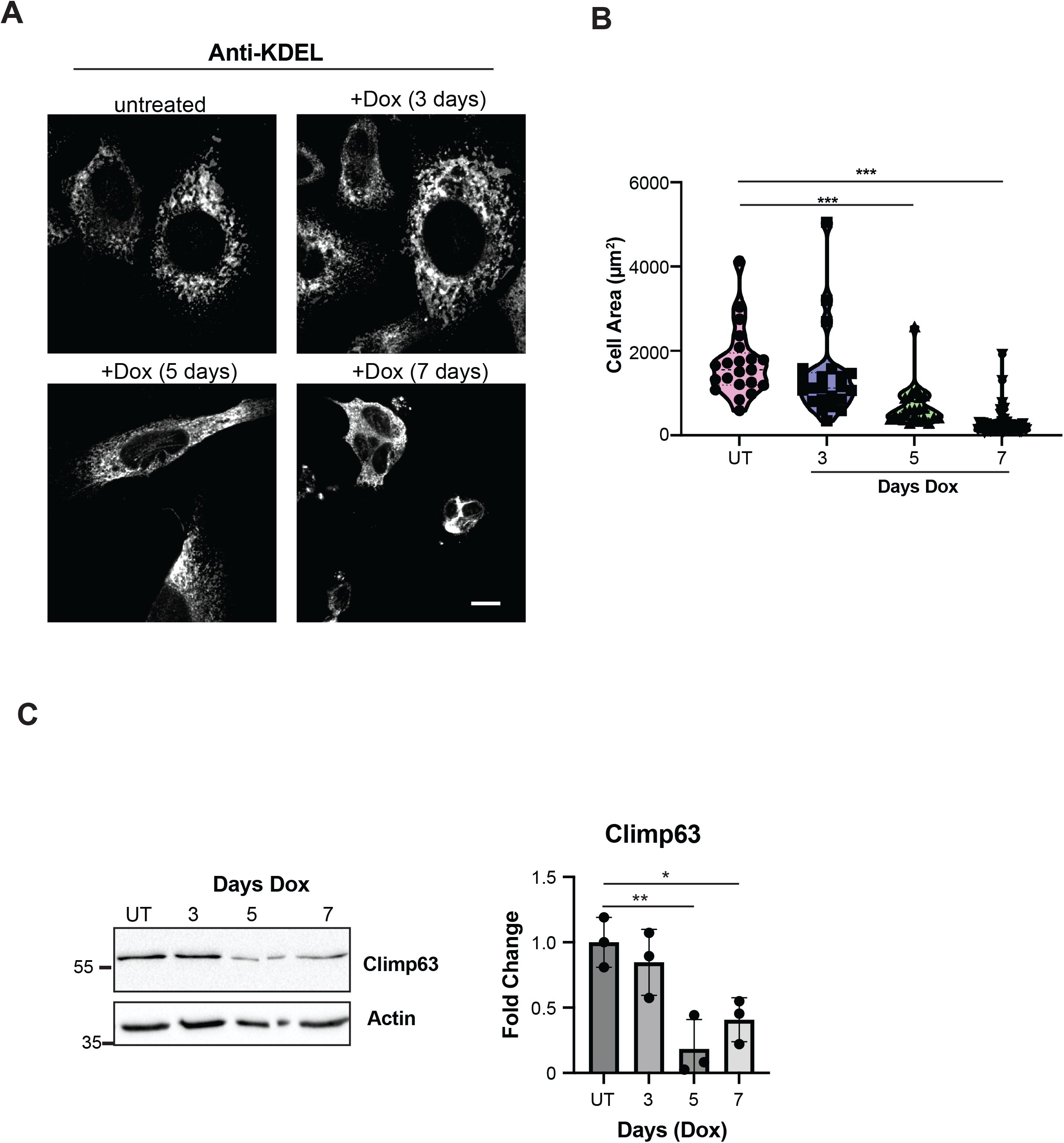
ER morphology changes in the absence of Grp170. **A**. E8 cells grown on coverslips were treated for the indicated times to induce Cre-lox editing of Grp170. Cells were stained for KDEL followed by anti-Rabbit 647 secondary to mark the ER. Scale bar in the bottom right panel is 10 µm (Bottom). **B.** Whole cell area was quantified using Nikon NIS-Elements software. Violin plots depicting the individual data points as well as the median, and first and third quartiles plotted *** p<0.0001. **C**. Protein lysates from the E8 clone treated with doxycycline for the indicated days was subject to western blot and probed with anti-Climp63 antibody. Data represent (n=3) the mean+/−SD; *p<0.05, **p<0.01.

## Discussion

In this study, we describe the isolation and characterization of a cell-based, inducible knock-out system to investigate the functional significance of Grp170, an ER resident molecular chaperone that exhibits both holdase and NEF activities. The importance of this chaperone was previously indicated by failed attempts to construct homozygous mouse knockouts (Kitao et al., 2001) and by the catastrophic effects on kidney function when Grp170 was depleted from nephrons in adult mice (Porter et al., 2022), but its requirement at the cellular level had not been determined. Using MEFs with regulated ablation of Grp170, we discovered that, other than BiP (Luo et al., 2006),(Paton et al., 2006), Grp170 is the only ER chaperone that is essential at the cellular level. Although Grp94 is required for mouse development, its depletion is tolerated in a number of tissues and cell lines (Wanderling et al., 2007). At the molecular level, murine Grp170 depletion disrupts several critical BiP functions including: (1) increased steady-state BiP substrate binding, consistent with defects in Grp170-dependent substrate release, (2) an ERAD defect, resulting in the accumulation of misfolded proteins in the ER, and (3) a pool of BiP becoming detergent insoluble, suggesting it becomes trapped with its aggregation-prone clients. In keeping with the hypothesis that loss of GRP impinges on BiP availability, BiP de-AMPylation, which provides a rapid restoration of functional BiP (Perera and Ron, 2023), was evident by day 3, when only ∼25% of Grp170 remained. Thus, its plausible that the cells are dying at later times, as Grp170 is further depleted, due to insufficient levels of BiP that are required to execute its essential functions.

Cell death may also arise due to unresolved UPR activation (Iurlaro and Muñoz-Pinedo, 2016). As the UPR transducers are regulated by BiP binding (Bertolotti et al., 2000),(Shen et al., 2002), this might also implicate diminished BiP availability as a cause of cell death. However, we observed irregularities in this stress response as Grp170 levels fell that might further diminish the UPRs ability to restore homeostasis. Importantly, the phosphorylation of eIF2α was sustained, and the resumption of translation was impaired as judged by decreased synthesis of NHK in the pulse-chase experiment, loss of Gadd34, which is the phosphatase responsible for dephosphorylation of eIF2α, and reduced expression of Climp63, a structural component of the ER. Another possibility for the lethality associated with loss of Grp170 is that an essential Grp170-dependent protein client(s) fails to mature properly in its absence. Currently, only a handful of proteins that transit the secretory pathway have been identified that directly interact with Grp170 (Lin et al., 1993),(Behnke and Hendershot, 2014), and at least some of them are likely aggregation-prone in the absence of Grp170 (Behnke et al., 2016). Overall, it is reasonable to assume that any or all of these effects contribute to the cellular requirement for Grp170.

In spite of our results revealing an essential role for Grp170 in MEFs, an unexpected finding was that over-expression of this chaperone was apparently not tolerated. In our single cell MEF clones there were multiple copies of the genomic region flanking the *Hyou1* gene due to the genetic instability associated SV40 transformation (Ray et al., 1990), but the *Hyou1* locus in these additional regions was deleted, resulting in only two alleles in each cell. In addition, although the bulk culture of the transformed MEFs expressed *Grp170* transcripts at twice the level of that in the untransformed MEFs, the protein levels were essentially the same, indicating that post-transcriptional mechanisms maintain wild-type levels of Grp170 protein. The reason for this tight regulation is not clear, but it is possible that excessive NEF activity might not be tolerated as it could affect BiP’s interactions with its clients. Further experiments are required to understand this result.

In contrast to the apparent essential nature of Grp170 in mammalian cells, the yeast homolog of GRP170, Lhs1, can be deleted and the cells continue to divide normally. However, similar to our observations in mammalian cells, loss of Lhs1 results in induction of the UPR. *Δlhs1* yeast also exhibit a mild temperature sensitive phenotype, a subtle defect in ER protein translocation (Craven et al., 1996), and consistent with the current study, *Δlhs1* yeast additionally exhibit an ERAD defect (Buck et al., 2013). Both higher eucaryotic cells and yeast possess a second ER localized NEF, Sil1, which is a functional homolog but lacks structural similarities with Grp170/Lhs1 and does not bind unfolded clients (Behnke et al., 2015). In yeast, loss of Sil1 induces a very mild UPR relative to loss of Lhs1 and no obvious protein processing defect, whereas deletion of both Lhs1 and Sil1 in combination is synthetic lethal (Tyson and Stirling, 2000). Interestingly, the *Δlhs1Δire1* double deletion is synthetically lethal in yeast but is rescued by Sil1 overexpression (de Keyzer et al., 2009). These data suggest that NEF activity is the essential function in yeast. Although we show that Sil1 levels increase when murine Grp170 is depleted (Figure 2D), these chaperones either exhibit independent essential functions in higher cells, or the level of Sil1 induction we observed was insufficient to rescue the loss-of-function phenotypes upon Grp170 ablation.

As opposed to the inability to produce a Grp170 null mouse (Kitao et al., 2001), the loss of Sil1 results in the production of viable progeny, although these mice phenocopy many of the symptoms associated with disease-associated mutations in SIL1 that result in Marinesco-Sjӧgren syndrome (Ichhaporia et al., 2018),(Zhao et al., 2005). Sil1 knockout in mice is also accompanied by UPR activation, leading to increased levels of Grp170 and revealing non-redundant aspects of these two ER NEFs. To avoid the catastrophic effects of complete Grp170 loss during development, we recently generated an inducible, kidney-specific Grp170 knockout mouse. Loss of Grp170 in the mouse kidney resulted in a dramatic phenotype and widespread kidney injury (Porter et al., 2022). As observed in the current study, there was strong induction of the UPR, alterations in ER morphology, and ultimately an apoptotic response. Although the cell-based, inducible knockout system reported here provides a more experimentally tractable system to understand how loss of Grp170 affects cell physiology, future work will better define how other components of the proteostasis network are affected upon the loss of Grp170. These data can then be compared to parallel ongoing efforts using our *in vivo* models to study Grp170 biology in other organ systems.

Another advantage of the new cell-based inducible Grp170 knockout system is that the phenotypic consequences of uncharacterized amino acid variants and polymorphisms in the gene encoding GPR170 on cellular physiology can now be studied. Consistent with a role of Grp170 in supporting the biogenesis of IgM antibodies (Lin et al., 1993), mutations in the *HYOU1* locus were identified in patients presenting with combined immunodeficiency as well as hypoglycermia (Haapaniemi et al., 2017). This latter symptom might be associated with the importance of GRP170 function during insulin maturation in the ER. Moreover, Grp170 limits aggregation of the Akita mutant form of insulin in mice by targeting this misfolded substrate for degradation via ER-phagy (Cunningham et al., 2019),(Cunningham et al., 2017). It is thus intriguing that polymorphisms in *HYOU1* are associated with diabetes in Pima Indians (Kovacs et al., 2002). GRP170, also known as ORP150, can be regulated independent of the UPR by proteasomal inhibition during hypoxic conditions (Zong et al., 2016). Recent studies have also implicated GRP170 in tumorigenesis associated with several cancers, including bladder, papillary thyroid, and breast. Of note, overexpression of GRP170 correlates with poor outcomes, increased cell mobility, and proliferation (Wang et al., 2023),(Wang et al., 2021),(Lee et al., 2021),(Hao et al., 2021). In addition, a requirement for GRP170 NEF activity in both cholera toxin and polyomavirus pathogenesis was uncovered, since GRP170 releases BiP from the toxin/virus, which is needed for ER to the cytosol retrotranlsocation and pathogenesis (Inoue and Tsai, 2015),(Williams et al., 2015). Finally, by using several models our group determined that Grp170 regulates the quality control and early biogenesis of the epithelial sodium channel (Buck et al., 2013), a major regulator of ion homeostasis and blood pressure (Porter et al., 2022)).

Overall, the findings reported here shed light on the essential cellular functions played by Grp170, including a major impact on BiP’s nucleotide-dependent activities, and provide a foundation for future studies to determine Grp170 substrate specificity, identify structure-function relationships in this dual-purpose chaperone, and define the defects associated with disease-causing alleles in the *HYOU1* locus (Haapaniemi et al., 2017).

## Materials and Methods

### Mice

LoxP sites were engineered in introns 1 and 24 of the *Grp170* (*Hyou1*) gene of C57Bl/6 mice to generate *Hyou^LoxP/LoxP^*mice, which were previously described (Porter et al., 2022). Both males and females *Hyou^LoxP/LoxP^*mice were fertile, as were their offspring. The mice were housed and treated in accordance with the Animal Use and Care Committee and the Animal Research Center at St. Jude Children’s Research Hospital, adhering to the National Institutes of Health guidelines. At weaning, DNA was isolated from mouse tail snips using DNeasy^®^ Blood & Tissue Kit (Qiagen, Valencia, CA) and genotyped. Primers for the wild-type allele and for the disrupted allele are listed below.

Oligos used for genotyping mice and MEF cells:

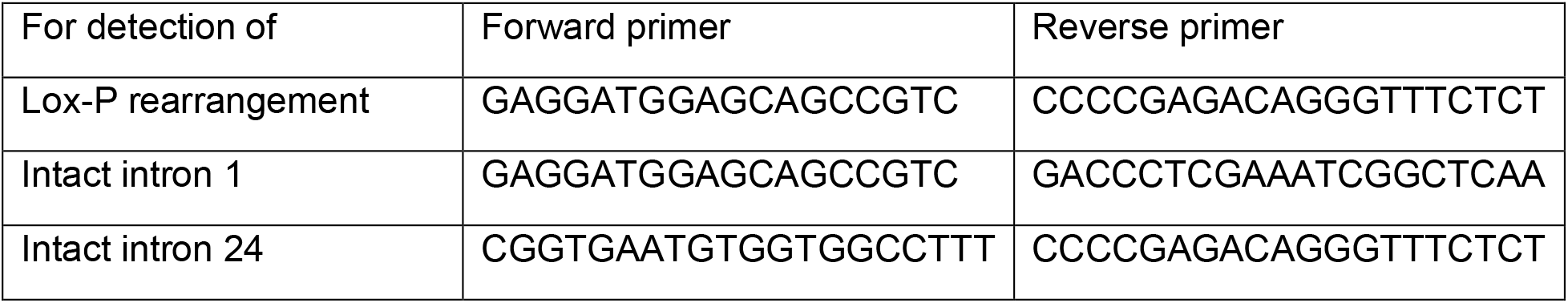

### Isolation of mouse embryonic fibroblasts from Hyou^LoxP/LoxP^ mice and transformation

Timed pregnancies were conducted using male and female *Hyou*^LoxP/LoxP^ (Porter et al., 2022). Embryos were surgically obtained at e12.5 day for the isolation of mouse embryonic fibroblasts (MEFs) using standard procedures (Kamijo et al., 1997). Briefly, organs from e12.5 embryos were removed and transferred to conical tubes containing DMEM/HEPES media (DMEM 4.5g/L glucose supplemented with 2mM L-Glutamine, Penicillin/Streptomycin, 0.1 mM non-essential amino acids, 5.5 μM 2-mercaptoethanol, and 10% fetal bovine serum). Organs were dispersed into single cells and plated on p100 dishes.

After 3 passages to select for fibroblasts, aliquots of *Grp170^LoxP/LoxP^* non-transformed MEFs were stored in liquid nitrogen., which are referred to as primary MEFs in this manuscript. To obtain transformed MEFs, the primary MEFs were plated, and at 80% confluency they were transfected with 4 μg of SV40 DNA per p60 dish using the Genecellin transfection reagent per manufacturer’s recommendations. Media was changed 24 hrs later. Cells were passaged to allow SV40-immortalized cells to outgrow non-transformed cells according to the Harding/Ron protocol (Harding et al., 2000). These *Grp170^LoxP/LoxP^/*SV40-transformed MEFs are referred to as parental MEFs.

### Cell Culture

MEF cells were maintained at 37°C and 8% CO_2_ in Dulbecco’s Modified Eagle’s Medium (DMEM) (Corning, 15013-CV) supplemented with 10% (v/v) Fetal Bovine Serum - TET Tested (Biotechne S10350), 55µM non-essential amino acids (Gibco 11140050), 1x 2-mercaptoethanol (Gibco 21985023), and 1% l-glutamine (Corning, 25-005-CI). To passage cells, TrypLE (Gibco) was used. 293T cells used for transient expression of the inducible Cre vector were maintained at 37°C and 5% CO_2_ in DMEM (Corning, 15013-CV) supplemented with 10% (v/v) fetal bovine serum (FBS) (Atlanta Biologicals S11550) and 1% l-glutamine (Corning, 25-005-CI). For all lines, cells were periodically tested for mycoplasma to ensure only mycoplasma-free cells were used.

### Creation of an inducible Cre construct and introduction into Grp170^LoxP/LoxP^ MEFs

The pSJL224 lentivirus backbone vector was obtained from the SJ vector core. This vector encodes a constitutive eGFP followed in-frame by the Tet-On 3G gene separated by a P2A ribosome cleavage site. The mCherry gene was introduced down-stream of the TRE 3G promoter, which was followed by a 2A peptide sequence and the Cre recombinase gene. This construct was designed to provide expression of both mCherry and Cre only in the presence of doxycycline (Supplemental Figure 1A). An initial test of this construct was performed by introducing it into two dishes of 293T cells using the Genecellin transfection reagent. Twenty-four hrs later, one dish was treated with 3 μg/ml of doxycycline in sterile water. The following day, GFP and mCherry expression were detected using FACS analysis (BD Biosciences Fortessa).

Lentiviruses were produced by combining standard 3^rd^ generation packaging and envelope plasmids with Cre-modified pSJL224 backbone vector in 0.25M calcium in sterile water. The transfection mixture was incubated at room temperature for 10-15 minutes and then added dropwise to HEK293T cells on a p100 dish. Cells were incubated at 37°C in 5% CO_2_ for 16-20 hours. Media was then aspirated, and 5ml of fresh DMEM was added. Twenty-four hours later, culture supernatants containing virus were harvested and passed through a 0.45 μM PVDF filter (Millipore, SLHV033RS). Viral titers were calculated and an MOI of 1 was used for transduction. The virus-containing media was mixed with DMEM/polybrene (final concentration 6 μg/mL) up to 5ml and was added to *Grp170^LoxP/LoxP^*MEFs on a p100 dish. Twenty-four hours later, an additional 5ml of DMEM was added to dilute lentiviral toxicity. After one week, a bulk culture of GFP^+^, mCherry^−^ cells was isolated by cell sorting (BD Biosciences Aria).

### Isolation of single cell clones with tight regulation of Cre

The bulk culture of GFP^+^ Hyou^LoxP/LoxP^ MEFs were resorted to obtain the brightest 15% GFP^+^ cells that were mCherry^−^ cells, and single cells were dispersed into 96-well plates. After expanding, 88 single cell clones were identified. An aliquot of each was treated with 3 μg/ml of doxycycline for 2 days, and the percent mCherry^+^ cells in each clone was determined by FACS analysis (Table 1). Cells were genotyped using the primers listed above.

### Fluorescence in situ hybridization

The Cytogenetic Shared Resource Laboratory at SJCRH designed a fluorescence *in situ* hybridization (FISH) assay to be used for this analysis. Purified *Hyou1* telomeric flanking fosmid DNA (WI1-2296B18) was used as a locus specific control. A 10kb subclone corresponding to the genomic *Hyou1* sequence between the two LoxP sites was PCR-amplified from the *Hyou1* fosmid WI1-1167J13 clone using an initial primer pair followed by a nested primer pair listed below. This probe was used identify the cre-lox programmed deletion. After a four-hour colcemid incubation, the cells were harvested using routine cytogenetic methods. The slides were pretreated with RNase A for 45 minutes prior to hybridization. After purification, the *Hyou1-*telomeric flanking fosmid DNA (WI1-2296B18) DNA was labeled with a red 580 dUTP (Enzo) by nick translation. The purified *Hyou1* 10kb genomic subclone was labeled with a green 496 dUTP (Enzo). The *Hyou1* (red) fosmid and the *Hyou1* (green) 10kb subclone probes were combined with sheared mouse cot DNA and hybridized to cells derived from the samples in a solution containing 50% formamide, 10% dextran sulfate, and 2X SSC. After overnight hybridization, the cells were washed and stained with DAPI (1μg/ml). Two hundred cells were analyzed for each of the experimental groups, and the numbers of intact vs deleted genes was determined. Cells in which the *Hyou1* gene is intact have paired red-green signals, while loci that have undergone the programmed deletion will have only the control flanking signals (red) present.

Primers used for isolating the genomic *Hyou1 probe*

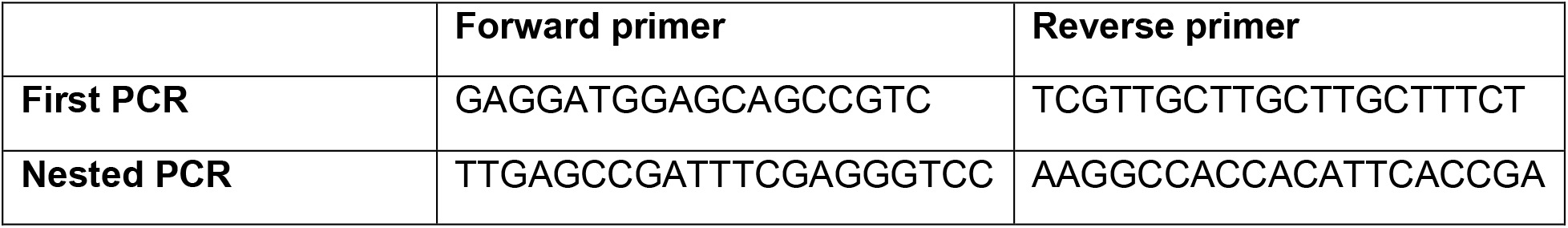

### RNA expression studies

Total RNA was isolated from MEF cell pellets using RNeasy® Plus Mini Kit (Qiagen), according to the company protocol. 2 μg total RNA was reverse transcribed using the High Capacity cDNA Reverse Transcription Kit (Applied Biosystems) and subjected to SYBR® Green chemistry-based qPCR using the QuantStudio 3 Real-time PCR System (Applied Biosystems, Foster City, CA). Primers were obtained from ThermoFisher Scientific. GAPDH was utilized as a reference gene control. Relative mRNA fold change was computed from the QuantStudio-generated Ct values by using the 2^−ΔΔCt^ method. Except in the case of Cre transcripts, mRNA values obtained from untreated cells were set to 100%, and values for the days after doxycycline addition were expressed relative to those in untreated cells. At least 3 biological replicates were performed, and each was assayed in triplicate. Statistical analyses were conducted using Prism 9 software package (GraphPad Software Inc). When comparing more than two groups a one-way ANOVA was performed with Tukey’s multiple comparisons test, where the mean of each group was compared to the mean of every other group.

### Oligos for mRNA expression

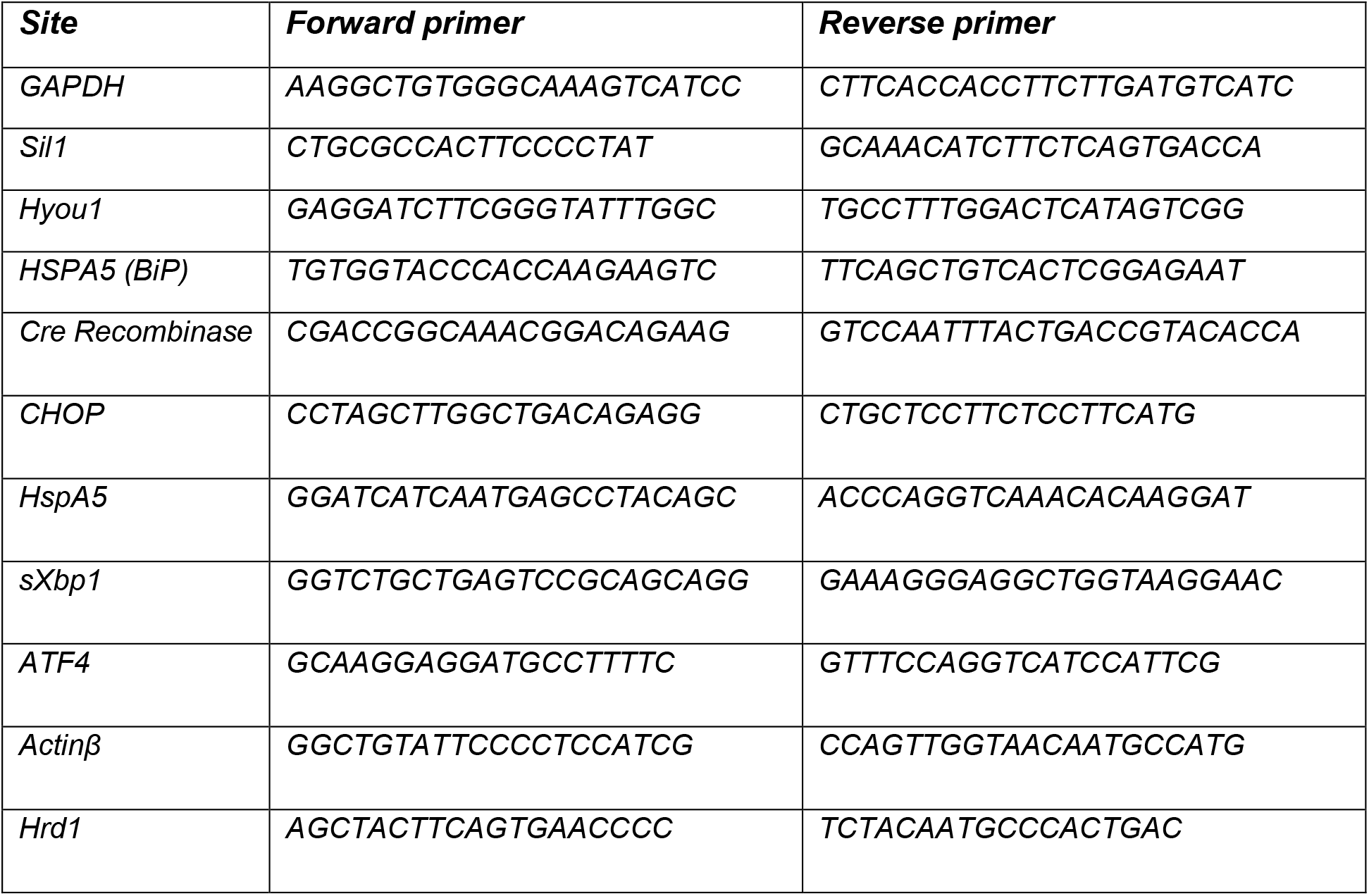

### Transfections, western blotting and immune reagents

E8 cells were plated with or without doxycycline for 3 day and then replated with the same conditions for 24 h before transfecting with the GeneCellin transfection reagent (BioCellChallenge, GC5000) according to the manufacturer’s instructions. Cell pellets were disrupted using NP-40 lysis buffer (0.5 M Tris-HCl pH 7.5, 0.15 M NaCl, 0.5% DOC, 0.5% NP40 substitute, 0.002% sodium azide) supplemented with 0.1 mM PMSF and 1x Roche complete protease inhibitor tablets and clarified by centrifugation for 15 min at maximum speed in a refrigerated microfuge to produce clarified cell lysates. Equivalent amounts of total protein, as determined by Coomassie Protein Assay Reagent (ThermoScientific, 23200) were loaded on gels. In the experiments to separate NP40 soluble and insoluble proteins the clarified lysate corresponded to the soluble material and the resulting pellet was solubilized by heating in SDS sample buffer at 95°C for 5 minutes followed by water bath sonication. The amount of insoluble pellet loaded was 6 times that of the corresponding soluble sample. In all cases, samples were electrophoresed under reducing conditions on SDS-polyacrylamide gels and transferred to PVDF membranes (Millipore #IPFL00010). Membranes were blocked with either 4% non-fat milk or fish gelatin (Sigma #G7041-100G) for 30 min at room temperature and incubated overnight in primary antibodies on an orbital rocker at 4°C. Membranes were serially washed in Tris buffered saline followed by Tris buffered saline with 0.1% Tween20 and incubated in secondary reagents for 1 hr at room temperature on an orbital rocker. Finally, membranes were serially washed again and imaged on a Li-Cor CLx imager and visualized using Image Studio™ (LI-COR Biosciences, Inc., Lincoln, NE) where fluorescent secondary antibodies were used. Alternatively, when HRP-conjugated secondary reagents were used the membranes were incubated in ECL Western Blotting Substrate (Pierce #32106) and developed on film (Denville Scientific #E3018) or imaged on a BioRAD Universal Hood II Imager and quantified using ImageJ 1.51h software (National Institutes of Health). Rabbit anti-rodent BiP (Hendershot et al., 1995), and GRP170 (Behnke et al., 2014) were previously produced in our lab. Immune serum specific for Cre (Cell Signaling, 12830S), CHOP (Cell Signaling #2895S) Hsc70 (Santa Cruz, SC-7298), eIF2α (#5324S) and p-eIF2α (Cell Signaling 9721L), PERK (Cell Signaling #3192), tubulin (Proteintech 66031), Cleaved-Caspase 3 (cell signaling #9661S), OS-9 (Abcam ab109510) and Climp63 (Bethyl Laboratories, A302-257A) were purchased from commercial venders. A monoclonal antibody specific for AMPylation and AMPylated recombinant BiP were kind gifts of the Itzen laboratory (Hamburg University). Secondary immune reagents were used at a dilution of 1:10,000 (anti-mouse; Cell Signaling #7076S) or anti-rabbit (#7074S). Values for protein signals were normalized to either Revert Total Protein Stain (Li-Cor, 926-11021), actin or tubulin and expressed relative to the untreated samples, which were set to 100%. For Cre protein, the day 2 value was set to 100%, and subsequent days of treatment were expressed relative to the day 2 value. At least 3 biological replicates were performed.

### Pulse-chase experiments

To measure the effects of Grp170 loss on an ERAD client, E8 cells were transfected with a vector encoding the NHK variant of α1AT that were either left untreated or incubated with doxycycline 4 days previously as described for NS1 experiments above. Cells were pre-incubated in 2 mL labeling media consisting of 1x DMEM without Cys and Met (Cys-/Met-) (Corning, 17-204CI) supplemented with 10% Tet free FBS dialyzed against PBS, 1% L-glutamine, and 1% antibiotic/antimycotic at 37°C for 30 min and then labeled with Express [35S] Labeling Mix (PerkinElmer, NEG072-007) for 15 min. After labeling, cells were put on ice, rinsed with PBS and 2ml of complete growth media supplemented with cold methionine and cysteine was added to the chase dishes, which were returned to 37°C and 8% CO_2_ for the indicated times. At the end of the chase period, cells were washed on ice 2x with PBS and then lysed with the NP40 buffer and clarified as described for western blotting. Samples were incubated with rabbit anti-α1-antitrypsin (A0409-1VL, Millipore Sigma) overnight at 4°C followed by Protein A agarose beads (CA-PRI-0100, Repligen) for 1 hr the following morning. Isolated protein was eluted with reducing Laemmli buffer by heating at 95°C for 5 minutes. Proteins were separated by SDS-PAGE and transferred to PVDF membrane as described for western blotting. Dried membranes were exposed to BAS storage phosphor screens (Cytiva, 28956475) and scanned using a phosphor imager (Typhoon FLA 9500 GE Healthcare). Signals were analyzed with the ImageQuant TL software (Cytiva) and expressed as percentage of the pulse (t=0) sample.

### Viability Assays

Cell Titer-Glo (Promega) viability assays were performed in 96-well plate according to the manufacturer’s instructions. Grp170 KO was induced by the addition of doxycyline (3 ug/ml) for the indicated number of days.

### Indirect Immunofluorescence

E8 cells grown on glass coverslips were treated with doxycycline for the indicated time points or were left untreated. Cells were then rinsed once with PBS, fixed for 20 min in 3.7% formaldehyde, rinsed with PBS containing 10 mM glycine (PBS-G), and permeabilized with 0.5% Triton X-100 in PBS-G for 3 min at room temperature. After washing, nonspecific binding sites were blocked by incubation for 5 min in PBS-G containing 0.25% (w/v) albumin. Coverslips were incubated for 1 hour with rabbit polyclonal anti-KDEL antibody (1:250 dilution;PA1-013;Thermofisher) to label the endoplasmic reticulum followed by washing in PBS-G. Cells were then incubated for 30 min with secondary antibody Alexa Fluor goat anti-rabbit 647 (1:500;A21245;Thermofisher). Following secondary antibody treatment, cells were washed extensively with PBS-G and mounted onto slides using Prolong Diamond Antifade Mountant with DAPI (P36962;Thermofisher). Confocal images were captured using a Nikon Eclimpse Ti2-E A1R inverted microscope outfitted with an oil immersion 100X objective (NA 1.45). Whole cell area was measured using the GFP signal using Nikon’s NIS-Elements software (Nikon Instrument, Melville, NY). Data were entered into Prism software (GraphPad, La Jolla, CA) for graphing and statistical analysis consisting of a Student’s t-test of the treatment groups vs the untreated control. Asterisks represent p<0.0001. The scale shown is 10 µm.

## Abbreviations

Grp170: glucose regulated protein-170r
BiP: binding protein
Grp94: glucose-regulated protein-94
Hsp70: heat shock protein-70
Hsp90: heat shock protein-90
ER: endoplasmic reticulum
NEF: nucleotide exchange factor
ERAD: endoplasmic reticulum associated degradation
UPR: unfolded protein response

## Acknowledgments

This work was supported by National Institutes of Health grants GM54068 (to L.M.H.) and the American Lebanese Syrian Associated Charities of St. Jude Children’s Research Hospital, NIH grants DK117162 (to T.M.B) and GM131732 and DK137329 (to J.L.B.). We also acknowledge Aidan Porter and the entire Brodsky and Hendershot labs for advice, reagents, and/or valuable insights during the course of this study. We are additionally indebted to the Itzen lab for providing a number of reagents for the AMPylation blots that were unavailable in the US and for sending a detailed protocol for using the antibody along with a positive control. We are also thankful for a suggestion by Dr. Joseph Chambers (University of Cambridge) to examine AMPylation of BiP and to Marcus Valentine for help with the FISH assay.

## Supplemental Figure Legends

**Supplemental Figure 1:** A doxycycline inducible system Grp170 KO MEF cell system. **A**. Schematic of a lentivirus backbone with a doxycycline-inducible Cre that can be readily monitored by fluorescence. **B**. 293T cells were transfected with the pSJL224 construct using transient expression protocols. 24 hrs after transfection, cells were left untreated or incubated with doxycycline for an additional 24 hrs and then analyzed by FACScan. GFP fluorescence reports on the presence of the construct, and mCherry denotes expression of doxycycline-inducible transcript that encodes mCherry and Cre recombinase. **C.** 293T cells were transfected with the pSJL224 construct or mock transfected and then cultured as in **B**. Cell lysates were prepared 24 hrs after transfection and analyzed by western blotting with anti-Cre.

**Supplemental Figure 2:** Confirmation of genetic editing and excision of *Hyou*1. **A**. DNA was isolated from the two single cell clones at the indicated times after incubation with doxycycline to induce Cre expression. RT-PCR was performed using two primer pairs that detect the intact gene as well as a pair to identify the recombined gene leading to excision of *Hyou1*. **B.** Schematic of the mouse *Hyou1* locus with notation of the placement of the primers used in **A** and the effects of Cre recombination on these sites. **C**. The number of *Hyou1* loci present in the two clones as well as their recombination were determined by FISH analyses of metaphase spreads. **D**. The levels of *Grp170* transcripts (black bars) and proteins (grey bars) were quantified in non-transformed MEFs, the MEFs after transformation with SV40, and in each of the two single cell clones. Each was expressed relative to the levels in the non-transformed MEFs. Three biological replicates were used to obtain the values shown.

**Supplemental Figure 3: The UPR is induced in the A10 clone.** RNA was isolated from A10 clones after Cre induction for the indicated number of days. qPCR was performed to detect targets of the UPR. Data represent (n=3) the mean+/−SD; *p<0.05, **p<0.01, ***p<0.001, ****p<0.0001.

**Supplemental Figure 4: Western blot quantitation for UPR targets. A**. Quantitation of western blot data presented in Figure 6B. Data represent the mean +/−SEM (n=3-5) *p<0.05, **p<0.01, ***p<0.001, ****p<0.0001. **B.** Quantitation of Gadd34 western blot from Figure 6C.

