## Supplementary figures and images for "Loss of Grp170 results in catastrophic disruption of endoplasmic reticulum functions"

### Supplemental Figures

Figure S1

A

pSJL224

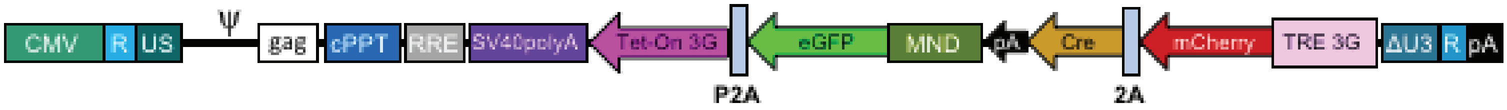


B

pSJL224

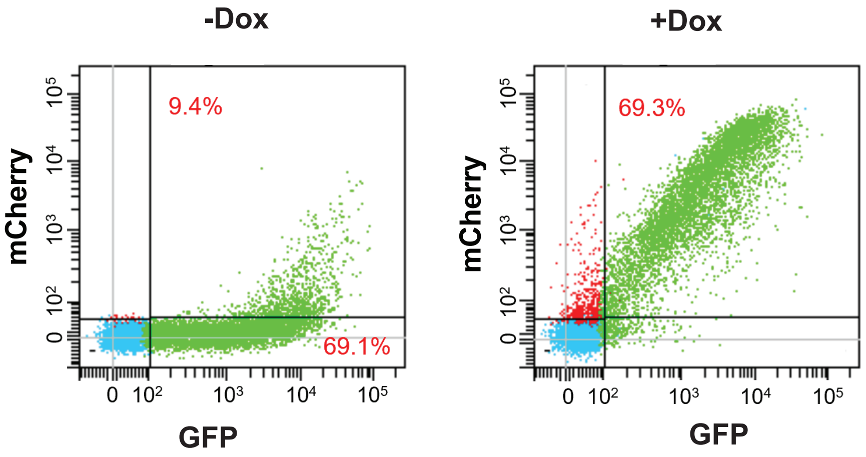


C

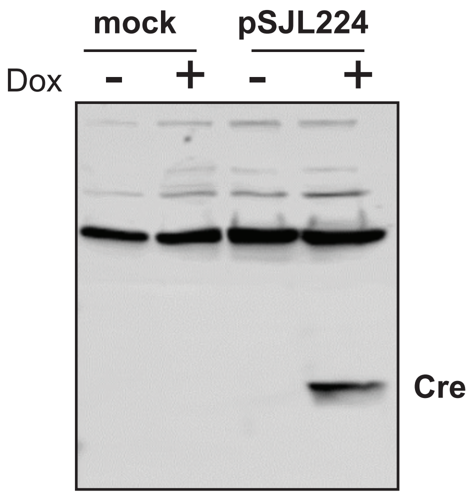

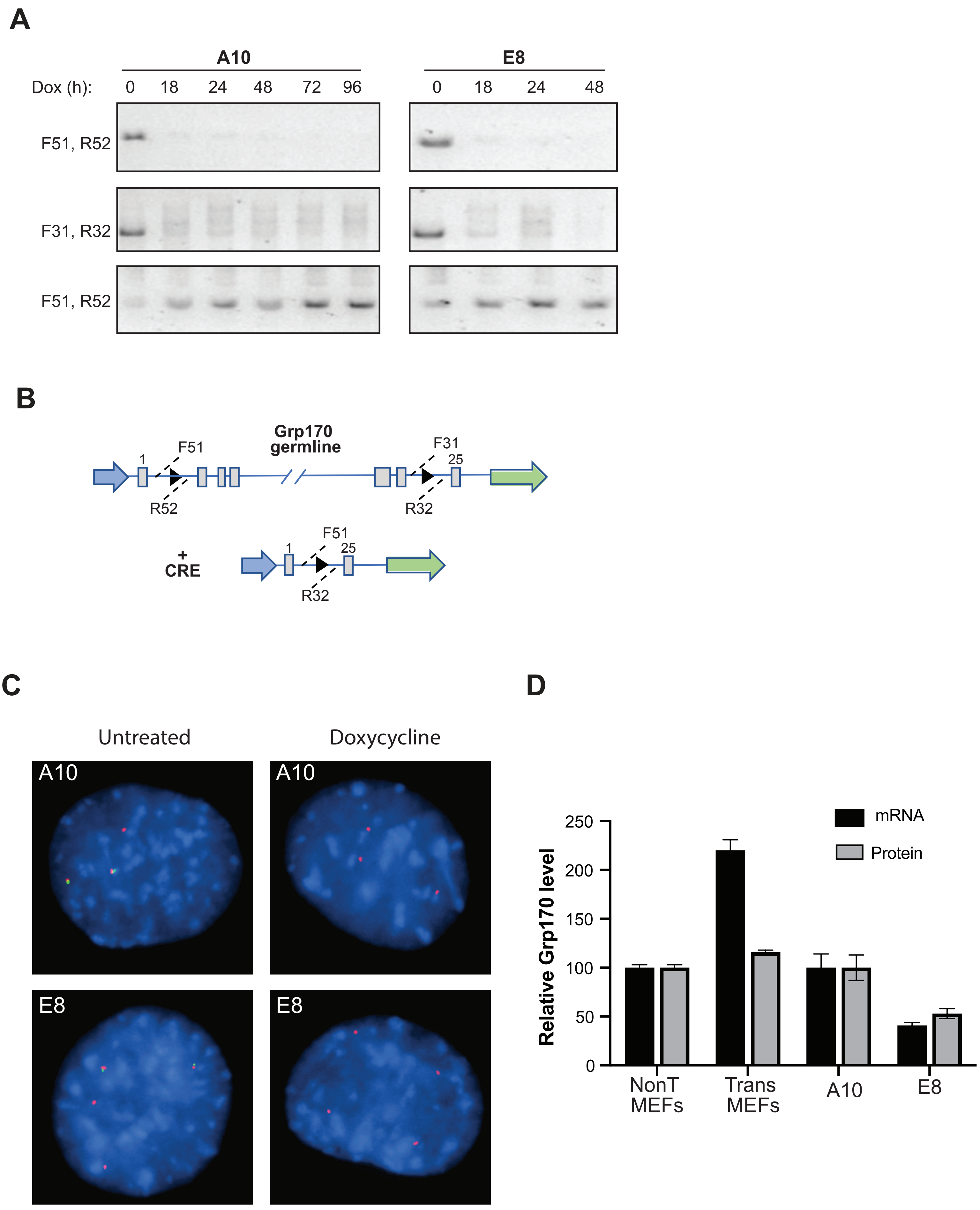

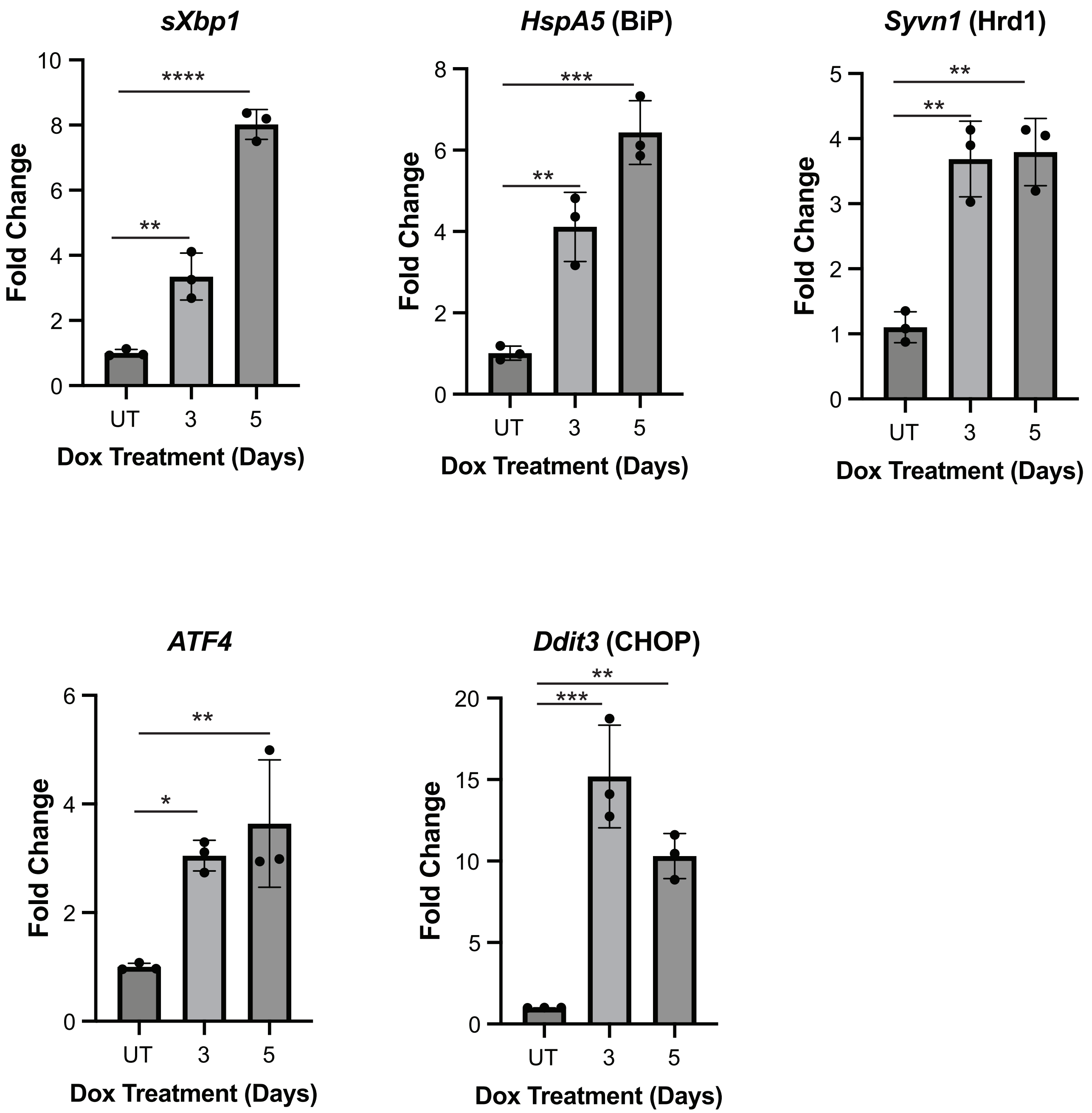

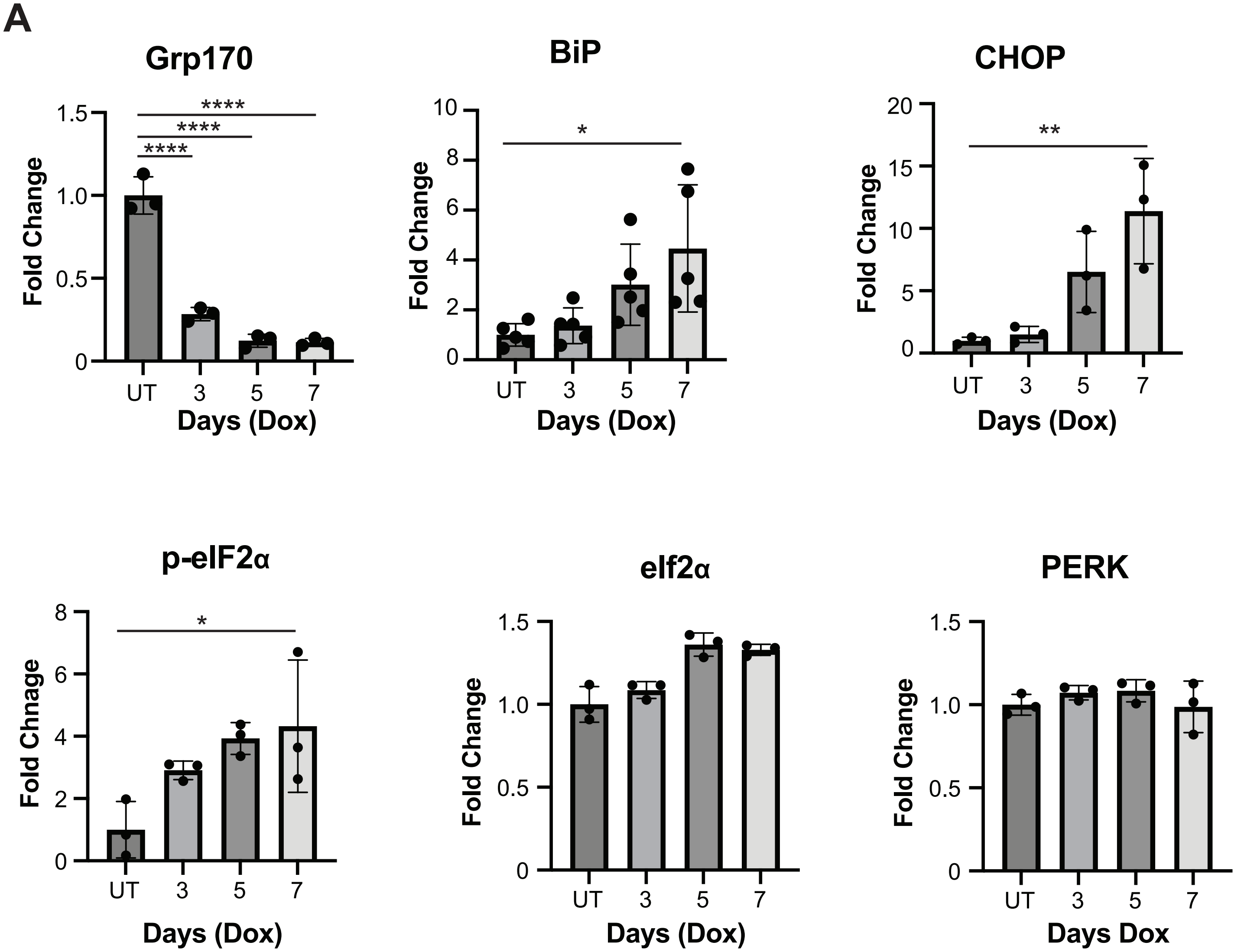

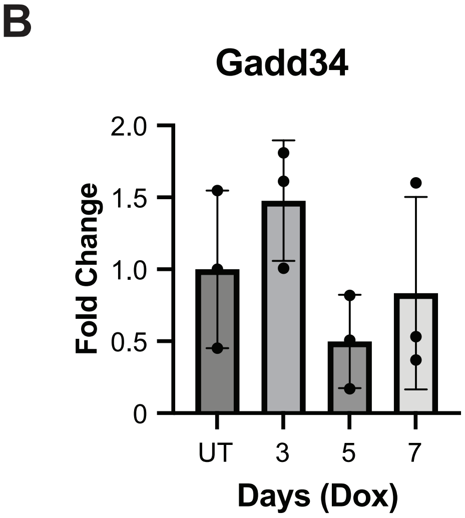
